# Super-rapid quantitation of the production of HIV-1 harboring a luminescent peptide tag

**DOI:** 10.1101/2020.04.10.033381

**Authors:** Seiya Ozono, Yanzhao Zhang, Minoru Tobiume, Satoshi Kishigami, Kenzo Tokunaga

## Abstract

In virological studies using HIV-1 proviral clones, virus production is normally monitored by either an RT assay or a p24 antigen capture ELISA. However, these assays are costly and time-consuming for routine handling of a large number of HIV-1 samples. In addition, sample dilution is always required in ELISA to determine p24 protein levels because of the very narrow range of detectable concentrations in this assay. Here, we establish a novel HIV-1 production assay system to solve the aforementioned problems by using a recently developed small peptide tag called HiBiT. First, we constructed a novel full-length proviral HIV-1 DNA clone and a lentiviral packaging vector in which the HiBiT tag was added to the C terminus of the integrase. Tagging the integrase with the HiBiT sequence did not impede the production or infectivity of the resultant viruses. Electron microscopy revealed normal morphology of the virus particles. Most importantly, by comparing between ELISA and the HiBiT luciferase assay, we successfully obtained an excellent linear correlation between p24 concentrations and HiBiT-based luciferase activity. Overall, we conclude that HiBiT-tagged viruses can replace the parental HIV-1 and lentiviral vectors, which enables us to perform a super-rapid, inexpensive, convenient, simple, and highly accurate quantitative assay for HIV-1 production.

## INTRODUCTION

The HIV-1 proviral DNA clone pNL4-3 (1) has been highly frequently used for *in vitro* HIV-1 studies worldwide for the last 34 years. Therefore, the resultant NL4-3 viruses are extraordinarily well characterized through a considerable amount of virological research, providing a solid knowledge base for data acquisition and/or interpretation of interexperimental data. Quantitation of viral supernatants from cells transfected with proviral DNA clones, including pNL4-3, is commonly and routinely performed by conducting a standard reverse transcriptase (RT) assay (2) or p24 antigen capture ELISA (3), both of which are time-consuming, taking up to 4-5 hours or more. Particularly in the case of commercially available ELISA kits for HIV-1 detection, the major disadvantages are the high cost of the assays, and an intrinsic limitation in the linear ranges that require extensive dilution. In this study, we constructed a novel pNL4-3-based full-length proviral HIV-1 DNA clone and a lentiviral vector with a small luminescent peptide tag called HiBiT. When this 11-amino-acid peptide tag binds to the larger counterpart protein called LgBiT, the full-length structure of the smallest luciferase protein called NanoLuc is formed (4) (Fig. 1A, upper panel). Due to its small size, HiBiT can be easily fused to the N or C terminus of the protein of interest. After expression of a HiBiT-tagged protein, the assay is performed by simply adding a lytic detection reagent containing LgBiT with the substrate, and the luminescence level of the HiBiT-tagged protein is quickly determined by a luminometer (Fig. 1A, lower panel). Using this system, we established a much faster (∼15 minutes), equally accurate and cheap method, comparable to the recently established SG-PERT assay (5, 6) (as described in Discussion), demonstrating that this molecular tool would largely provide a cost/time-effective routine quantification of HIV-1 and lentiviral vectors.

**Figure 1.**
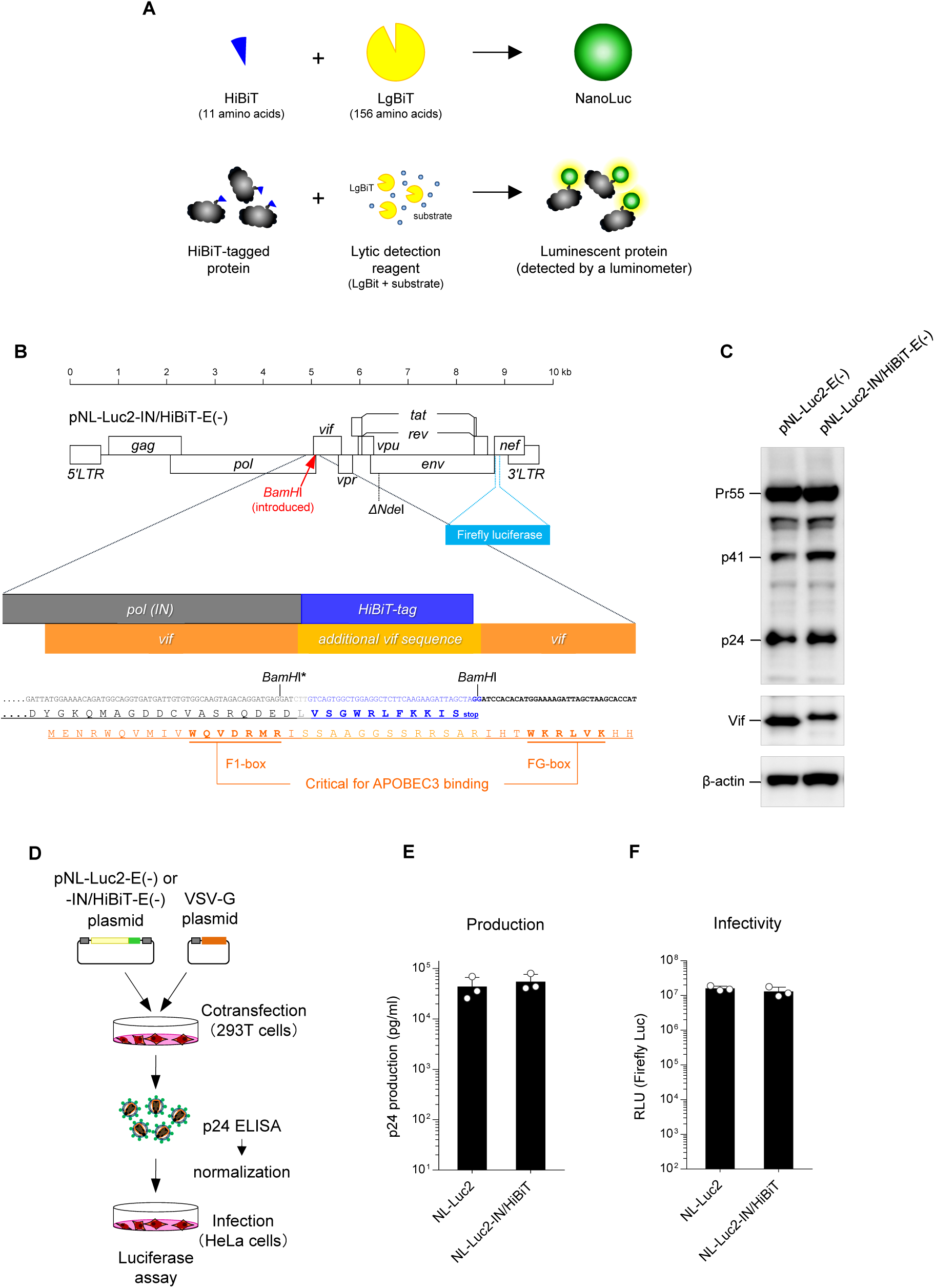
Luciferase reporter HIV-1 proviral DNA with a HiBiT tag maintains the expected levels of production and infectivity. (A) Schematic diagram of the HiBiT luciferase assay system (adapted from Promega’s website; https://www.promega.com/products/protein-detection/protein-quantification/nano-glo-hibit-lytic-detection-system/?catnum=n3030&catNum=N3030). (B) Construction of the HiBiT-tagged HIV-1 proviral DNA designated pNL-Luc2-IN/HiBiT-E^-^. A HiBiT tag (11-amino acid sequence shown in blue) was inserted into the C terminus of the integrase of pNL-Luc2-E^-^R^+^ in which *BamH*I was newly introduced (the upstream *BamH*I site with an asterisk was disrupted by inserting the HiBiT tag). The amino acid sequence shown in light orange represents the *vif* gene with an unrelated sequence created by the HiBiT tag insertion. (C) Western blot analysis performed by using extracts from 293T cells transfected with either pNL-Luc2-E-R+ or pNL-Luc2-IN/HiBiT-E-. Antibodies specific for p24 (upper), Vif (middle), and β-actin (lower) were used. (D) Schematic flowchart of the experimental procedure for HIV-1 virion production and infectivity assays. 293T cells were transfected with either pNL-Luc2-E-R+ or pNL-Luc2-INtc together with pC-VSVg, and 48 hours later, the viral supernatants were harvested and subjected to HIV-1 p24 ELISA to determine the level of virion production. Transfection efficiencies were normalized to the activity of firefly luciferase. Equivalent amounts of p24 antigen of VSV-G-pseudotyped viruses were used for the infection of HeLa cells. After 48 hours, cells were lysed, and firefly luciferase activities were measured to determine viral infectivity. (E, F) Comparison of virion production (*E*) and viral infectivity (*F*) of NL-Luc2 and NL-Luc2-IN/HiBiT viruses (mean ± S.D. from three independent experiments).

## RESULTS

### Viruses carrying the HiBiT-tagged integrase retain the wild-type level of viral production and infectivity

To generate a HiBiT-tagged HIV-1 proviral clone, we first introduced a restriction enzyme site of *BamH*I at the C-terminal end of the integrase (IN) open reading frame (ORF) of the HIV-1 luciferase reporter virus construct pNL-Luc2-E(-)R(+), and used this as a cassette for insertion of the HiBiT tag (Fig. 1B). Then, we transfected 293T cells with the resultant construct pNL-Luc2-IN/HiBiT-E(-) or the parental construct pNL-Luc2-E(-), subjected the cell lysates to Western blotting analyses, and found that the HiBiT tag sequence did not affect the expression of the HIV-1 Gag and Vif proteins (Fig. 1C). Then, this proviral clone was cotransfected into 293T cells with the VSV-G expression plasmid, and the levels of p24 antigen in the viral supernatants were measured by ELISA (Fig. 1D). To normalize the transfection efficiency, transfected cells were lysed and subjected to assays for firefly luciferase activity. The luciferase reporter virus carrying the HiBiT-tagged IN (Luc2-IN/HiBiT virus) showed a wild-type (WT) level of virion production (Fig. 1E). Equal amounts of VSV-G-pseudotyped Luc2-WT and IN/HiBiT viruses were inoculated into 293T cells, and luciferase activities were measured to determine viral infectivity (Fig. 1D). The Luc2-IN/HiBiT virus was found to be equally infectious as the WT virus (Fig. 1F). Thus, tagging the C-terminal end of the IN ORF with the HiBiT sequence does not interfere with the productivity or infectivity of HIV-1.

### Insertion of the N-terminal *vif* sequence into the C terminus of IN confers intact anti-A3G activity to the resultant virus, leading to efficient replication in CD4-positive T-cells

To enable us to perform replication assays in primary cells that express the restriction factor APOBEC3G (A3G), the intact function of Vif in the Luc2-IN/HiBiT virus needed to be verified, since an unrelated amino acid sequence derived from the HiBiT tag was inserted in the N-terminal sequence of Vif that overlaps with the C terminus of the IN ORF (although critical domains of Vif for APOBEC3 binding (7-10) were unchanged) (Fig. 1A). To test whether anti-A3G activity remained intact in the Vif protein of the Luc2-IN/HiBiT virus, we cotransfected 293T cells with the VSV-G expression plasmid and either the parental pNL-Luc2, Vif-deficient pNL-Luc2-F(-), or pNL-Luc2-IN/HiBiT-E(-) construct, together with either a control plasmid or a plasmid encoding A3G, and used viruses obtained from the transfected cells for infection. A3G expression resulted in a marked reduction in infectivity of the Luc2-IN/HiBiT virus to the level of the NL-Luc2-F(-) virus (Fig. 2C), indicating that the Vif protein of the Luc2-IN/HiBiT virus lacks anti-A3G activity. To rescue the lost function of Vif whose sequence was interrupted by the HiBiT tag, we first disrupted the initiation codon of Vif overlapping with the C terminus of the IN ORF. After introducing the BlpI site, we inserted an oligo DNA linker corresponding to an N-terminal *vif* sequence immediately downstream of the stop codon of the HiBiT tag and codon-optimized the C-terminus of the IN (to avoid homologous recombination during reverse transcription) (Fig. 2A). We prepared cell lysates from 293T cells transfected with this construct designated pNL-Luc2-IN/HiBiT-E(-)Fin, performed Western blot analyses and confirmed the expression of Vif protein at the correct size (Fig. 2B). The Luc2-IN/HiBiT-Fin virus produced from cells transfected as described above was tested for infectivity and was found to be fully resistant against A3G (Fig. 2C), demonstrating that the resultant virus gained intact Vif activity, as expected.

**Figure 2.**
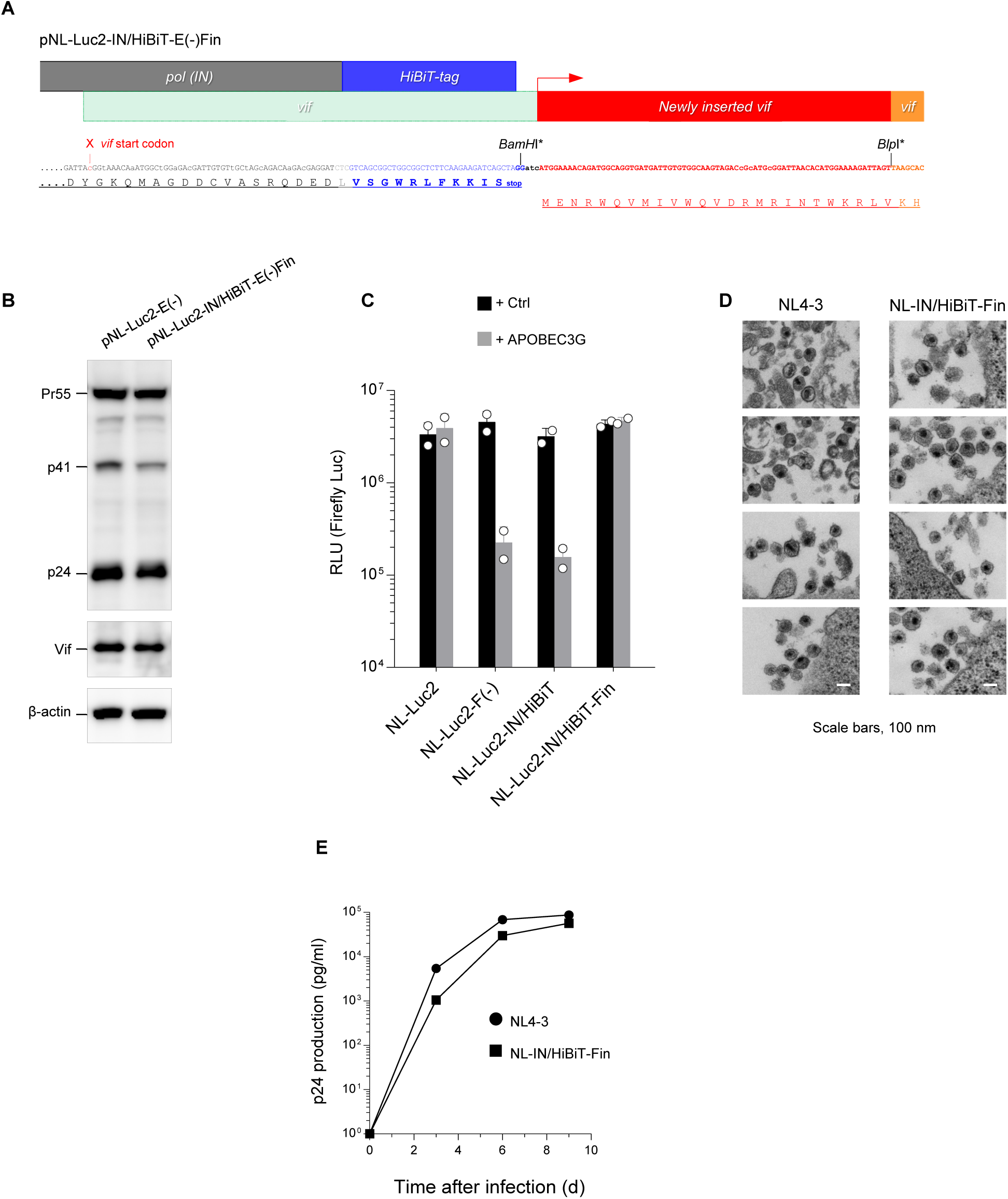
Rescued Vif activity results in intact anti-APOBEC3G activity and replication ability. (A) Reconstruction of the HiBiT-tagged HIV-1 proviral DNA that rescues Vif function. After mutating the *vif* initiation codon (indicated by the red arrowhead), the fragment encoding the 25-amino acid sequence (shown in red) was inserted immediately downstream of the HiBiT tag into *BamH*I and *Blp*I sites, the latter of which was newly introduced at the N terminus of the Vif sequence (both sites with asterisks were disrupted by inserting the fragment). (B) Anti-p24 (upper), anti-Vif (middle), and anti-β-actin (lower) immunoblots of cell extracts transfected with either pNL-Luc2-E^-^R^+^ or pNL-Luc2-IN/HiBiT-E(-)Fin. (C) Infection of MAGIC5 cells by HIV-1 Env-pseudotyped luc-reporter WT (NL-Luc2), Vif-deficient NL-Luc2 (NL-Luc2-F(-)), HiBiT-tagged NL-Luc2 (NL-Luc2-IN/HiBiT), and HiBiT-tagged/Vif-inserted NL-Luc2 NL-Luc2-IN/HiBiT-Fin), produced from cells expressing a vector control (black) or APOBEC3G (gray). Data from two experiments are shown (mean ± S.D., n = 3 technical replicates. (D) A representative transmission electron microscopy image of NL4-3 and NL-IN/HiBiT-Fin virions accumulated at the surface. Bars, 0.1 μm. (E) Multiple rounds of virus replication in peripheral blood mononuclear cells. PHA-IL-2-stimulated cells (2.5 x 10^5^) were infected with 25 ng of p24 antigen of either the NL4–3 or NL-IN/HiBiT-Fin virus. Supernatants were harvested at the indicated times, and virus replication was monitored using p24 ELISA. The data shown are representative of three independent experiments (mean ± S.D. of three technical replicates). d, days.

To determine whether an Env-intact full-length version of the viruses could normally replicate in CD4-positive T-cells, we generated a pNL4-3-based IN/HiBiT-Fin proviral clone (pNL-IN/HiBiT-Fin). We first performed transmission electron microscopy analyses using M8166 cells infected with either the resultant IN/HiBiT-Fin virus or the parental NL4-3 virus. Electron microscopic images revealed that mature particles of the IN/HiBiT-Fin virus were morphologically normal and indistinguishable from those of the parental NL4-3 virus, demonstrating that the insertion of the HiBiT tag has no influence on virion morphogenesis (Fig. 2D). Then, we infected PHA-stimulated primary CD4-positive T-cells with the WT or IN/HiBiT-Fin virus. Then, we monitored virus replication by measuring the levels of p24 production in the supernatants and found that the replication kinetics of the IN/HiBiT-Fin virus was only slightly delayed compared with that of the parental virus, confirming that the IN/HiBiT-Fin virus is replication-competent (Fig. 2E). Taken together, the results indicate that the addition of the N-terminal *vif* sequence into the C terminus of IN confers anti-A3G activity, resulting in efficient viral replication in primary CD4-positive T-cells.

### HiBiT-mediated luciferase activity strongly correlates with p24 concentrations in viral supernatants

Finally, we assessed whether HiBiT-tagged viruses could be normally detected by HiBiT-based luciferase assays and whether a correlation between HiBiT-mediated luciferase activity and the p24 concentration could be observed. We prepared a high-titer virus stock by propagating the IN/HiBiT-Fin virus produced by M8166 cells and measured p24 by a commercially available ELISA kit. Then, we serially diluted the samples as standard viruses in 10-fold increments, and subjected them to HiBiT luciferase assays as follows: we added equal amounts of samples and reagents, mixed them well, incubated them for 10 minutes, and performed measurements for 1 to 10 seconds per well by using a luminometer (Fig. 3A). HiBiT-mediated luciferase activity strongly correlated with p24 values in viral supernatants diluted on the order of μg (Fig. 3B), as well as highly diluted samples on the order of pg (Fig. 3C), with an excellent correlation between p24 levels and HiBiT luciferase activity (R^2^=1 and R^2^=0.9994, respectively). This indicates that the HiBiT activities can be directly translated to p24 antigen levels, routinely by this quick and easy method; i.e., we prepared the viral supernatants with known levels of p24 antigen and serially diluted them, and we could readily create a standard curve of HiBiT luciferase activity to determine the p24 levels of the samples of interest by measuring the activity in this assay. Overall, we conclude that the IN/HiBiT-Fin virus, in which the HiBiT tag and Vif linker were inserted into the C terminus of IN, is completely analogous to the parental HIV-1 strain and will be a highly convenient research tool for future HIV-1 studies.

**Figure 3.**
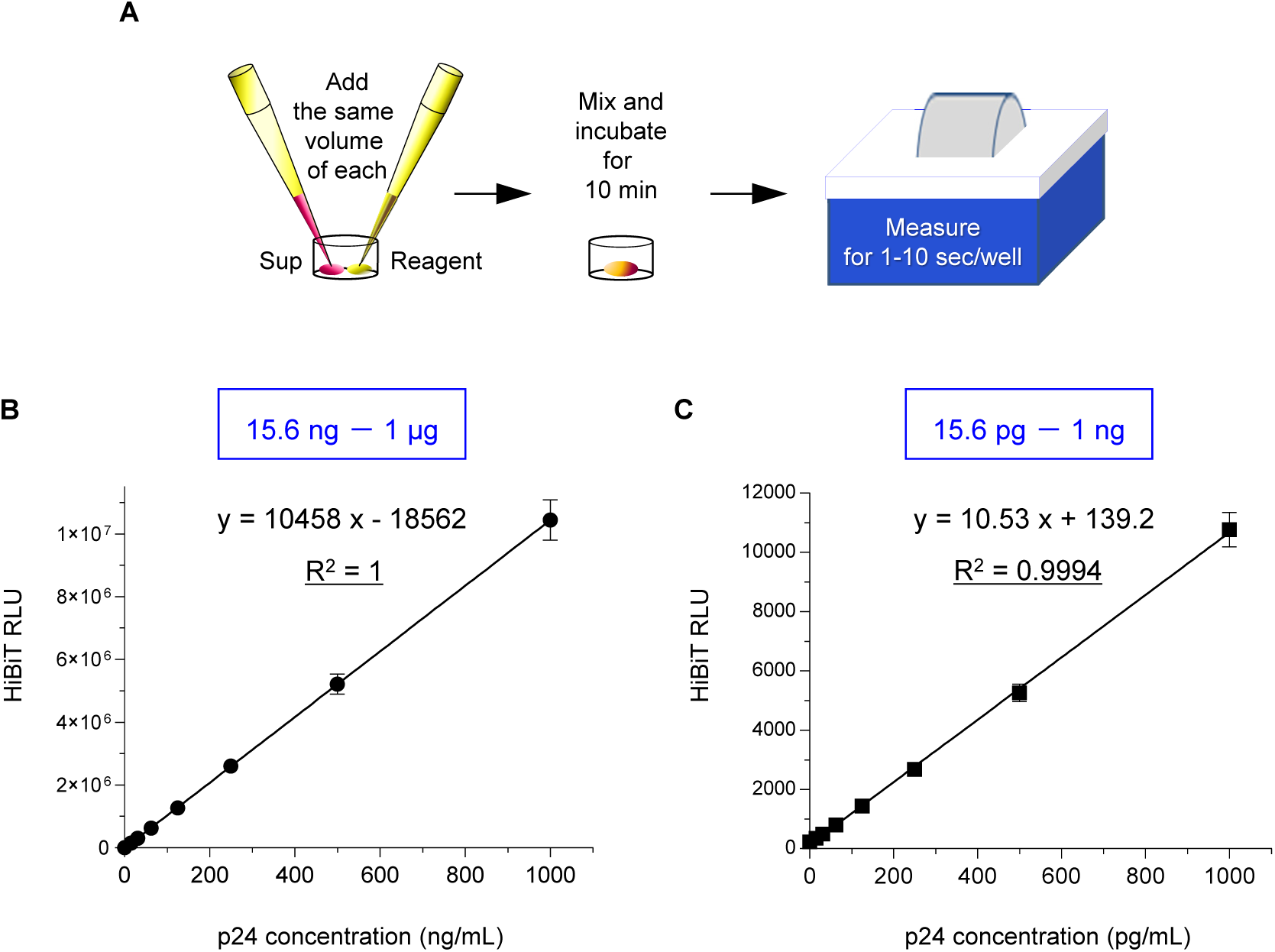
HiBiT-based luciferase assays show a strong correlation with p24 values in viral supernatants. (A) Schematic flowchart of the experimental procedure for HiBiT-based HIV-1 virion production assays. (B, C) Correlation between known levels of p24 antigen (*B*, high doses, 15.6 ng to 1 μg; *C*, low doses, 15.6 pg to 1 ng) measured with p24 antigen ELISA and HiBiT-derived luciferase activity measured with luciferase assays by using supernatants containing replication-competent NL-IN/HiBiT-Fin virus prepared from a human CD4-positive T-cell line. The equation and the coefficient of determination (R^2^) are shown as indicators of the linearity of this relationship (mean ± S.D. from at least three independent experiments).

## DISCUSSION

In this study, we established a novel system for the highly accurate, convenient, and super-rapid detection of the well-analyzed HIV-1 strain pNL4-3 carrying a luminescent peptide-tagged IN. This method provides an efficient routine for quantitating the virus production of full-length infectious HIV-1. Several research groups have developed different real-time RT-PCR methods for virion quantitation of lentiviral vectors or whole HIV-1 viruses, which target various HIV-1 viral genomes (reviewed in (11)) and transgene mRNA (12). These methods require RNA extraction and DNase treatment steps as well as a real-time RT-PCR procedure, taking a total of more than 4-5 hours. More recently, Vermeire *et al*. reported a real-time PCR-based, accurate, fast and relatively cheap method for retroviral quantification designated SG-PERT (SYBR Green I-based PCR-enhanced reverse transcriptase), directly determining RT activity in viral supernatants (5). According to the report, the reagent cost of the SG-PERT assay is approximately 10 times lower per retroviral quantification compared with the measurement of p24 antigen levels with a commercial ELISA kit, and the assay itself only requires less than 2 hours of hands-on time. In our study, the cost of the HiBiT-based assay reagents per well to quantify viral supernatants is almost equivalent to that of the SG-PERT assay. In addition, due to the wide range of detectable HiBiT activity, this assay does not require sample dilution similar to the SG-PERT assay, whereas p24 ELISA frequently requires three different dilutions that make the assay more time-consuming. Notably, whereas the whole ELISA procedure normally takes 4-5 hours (including sample preparation, incubation, and washing), that of the HiBiT-based assay (a one-step procedure) only takes up to ∼15 minutes, which is even faster than the aforementioned SG-PERT assay duration.

One clear limitation of this study needs to be mentioned: our system cannot be applied for bulk virus samples directly obtained from patients unless they are molecularly cloned and reconstructed into HiBiT-harboring proviral DNAs. Therefore, this method is specific for molecular clone-based virological studies. By taking advantage of this technique, we also constructed a lentiviral packaging vector carrying the HiBiT tag sequence, which is also fused to the C terminus of the integrase gene (Fig. 4A), since this could make lentiviral work more convenient. Consistent with the aforementioned observation using the proviral DNA, its virion production and infectivity were comparable to those of the WT lentiviral vector (Figs. 4B and 4C).

**Figure 4.**
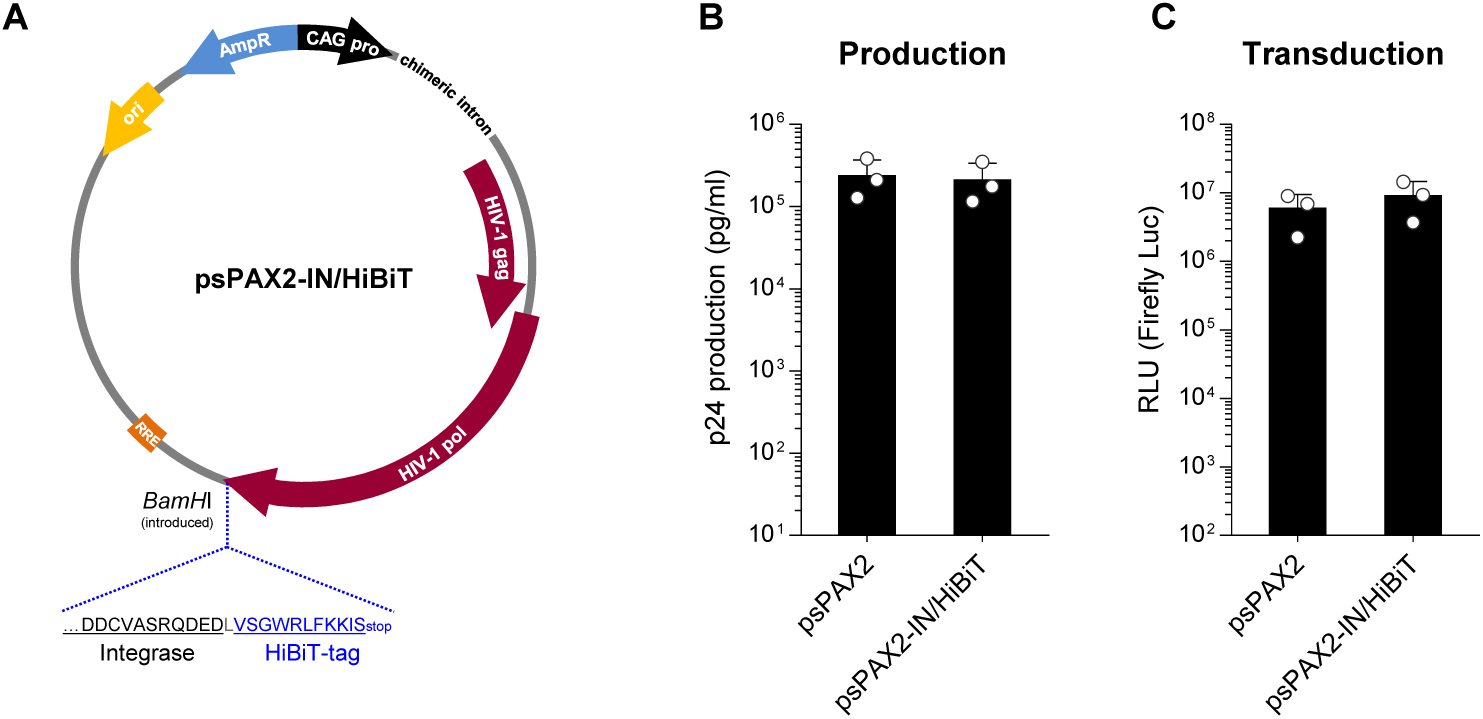
The lentiviral system using a lentiviral packaging vector carrying the HiBiT tag sequence exhibits WT levels of virion production and infectivity. (A) Construction of the HiBiT-tagged lentiviral packaging vector designated psPAX2-IN/HiBiT. A HiBiT tag was inserted into the C-terminus of the integrase of psPAX2 in which *BamH*I was newly introduced. (B, C) Comparison of lentiviral production (*B*) and transduction (*C*) resulting from lentiviruses produced from cells transfected with the luciferase-expressing transfer vector pWPI-Luc2 and VSV-G expression plasmid together with either the parental psPAX2 or the psPAX2-IN/HiBiT lentiviral packaging vector (mean ± S.D. from three independent experiments).

Thus, our HiBiT-based assay offers the following advantages: 1) super-rapid (∼15 minutes); 2) inexpensive (∼ 0.5 USD/sample); 3) practical and convenient (no dilution required); 4) extremely simple (one-step procedure); and 5) highly accurate (linear scale over 6 orders of magnitude ranging from 10 pg to 1 μg of p24 antigen). Overall, we conclude that the HiBiT-tagged proviral HIV-1 DNA and lentiviral vector can be used to replace their parental vectors, making them powerful and reliable molecular tools for future HIV-1 studies.

## METHODS

### DNA construction

The vesicular stomatitis virus glycoprotein (VSV-G) expression vector pC-VSVg, the HIV-1 proviral construct pNL4-3, its indicator construct pNL-Luc2-E^-^R^+^ and the A3G expression plasmid pC-hA3G-HA have been described previously (13, 14). To generate a luciferase reporter HIV-1 proviral DNA carrying HiBiT tag, the *Sbf*I/*EcoR*I fragment of pNL-Luc2-E^-^R^+^ was subcloned into *Pst*I/*EcoR*I-digested pUC19 (termed pUNL-PE), and the *BamH*I site was introduced by QuikChange mutagenesis (Stratagene) into pUNL-PE using the following specific oligonucleotides (restriction enzyme sites underlined); 5’-GAC AGG ATG A<u>GG ATC C</u>AC ACA TGG AAA AG-3’ and its antisense. The resultant plasmid was digested with *BamH*I, and used for the insertion of an oligonucleotide linker corresponding to HiBiT-tag (5’-GAT CTT GTC AGT GGC TGG AGG CTC TTC AAG AAG ATT AGC TAG-3’ and 5’-GAT CCT AGC TAA TCT TCT TGA AGA GCC TCC AGC CAC TGA CAA-3’). The *Pst*I/*EcoR*I fragment of the resultant vector pUNL-PE-IN/HiBiT was cloned back into *Sbf*I/*EcoR*I-digested pNL-Luc2-E^-^R^+^, and the generated plasmid was designated pNL-Luc2-E^-^-IN/HiBiT. The Vif-fixed version of IN/HiBiT virus construct was created as follows. The initiation codon of the *vif* gene was mutated, and the C terminus of the IN was codon-optimized by QuikChange mutagenesis in pUNL-PE-IN/HiBiT using the following specific oligonucleotides: 5’-CAT CAG GGA TTA CGG TAA ACA AAT GGC TGG AGA CGA TTG TGT TGC TAG CAG ACA AGA CGA GGA TCT-3’ and its antisense (the resultant construct was designated pUNL-PE-IN/HiBiT-ΔF). The *Blp*I site was introduced by QuikChange mutagenesis into pUNL-PE-IN/HiBiT-ΔF using the following specific oligonucleotides (restriction enzyme sites underlined): 5’-GAA AAG ATT A<u>GC TAA GC</u>A CCA TAT G-3’ and its antisense. The resultant plasmid was digested with *BamH*I and used to insert an oligonucleotide linker encoding the N-terminal Vif sequence (5’-GAT CAT GGA AAA CAG ATG GCA GGT GAT GAT TGT GTG GCA AGT AGA CCG CAT GCG GAT TAA CAC ATG GAA AAG ATT AGT-3’ and 5’-TTA ACT AAT CTT TTC CAT GTG TTA ATC CGC ATG CGG TCT ACT TGC CAC ACA ATC ATC ACC TGC CAT CTG TTT TCC AT-3’) downstream of the HiBiT-tag sequence. The *Pst*I/*EcoR*I fragment of the resultant vector pUNL-PE-IN/HiBiT-Fin was cloned back into *Sbf*I/*EcoR*I-digested pNL-Luc2-E-R+ or pNL4-3, and the generated plasmids were designated pNL-Luc2-E^-^-IN/HiBiT-Fin or pNL-IN/HiBiT-Fin, respectively. To create a lentiviral packaging vector carrying HiBiT-tag, the *Sbf*I/*Sac*I fragment of psPAX2 (15) was subcloned into *Pst*I/*Sac*I-digested pBluescript II SK(+) (termed pBSPAX-PS), and the *BamH*I site was introduced by QuikChange mutagenesis into pBSPAX-PS using the following specific oligonucleotides (restriction enzyme sites underlined); 5’-GGA TGA <u>GGA TCC</u> ACA CAT G −3 and its antisense. The resultant plasmid was digested with *BamH*I, and used for the insertion of an oligonucleotide linker corresponding to the aforementioned HiBiT-tag. The *Pst*I/*Sac*I fragment of the resultant vector pBSPAX-IN/HiBiT was cloned back into *Sbf*I/*Sac*I-digested psPAX2, and the generated plasmid was designated psPAX2-IN/HiBiT. To create a lentiviral transfer vector expressing firefly luciferase, the *Sal*I/*Sal*I fragment of pWPI (15) was subcloned into *Sal*I-digested pcDNA3.1 (Invitrogen). The resultant plasmid was digested with *EcoR*I to remove the internal ribosome entry site and EGFP, and further digested with *Pme*I to insert a PCR-amplified/*Pme*I-digested codon-optimized luciferase (Luc2) fragment. Then, the *Sal*I/*Sal*I fragment of the plasmid was cloned back into *Sal*I-digested pWPI, and the generated plasmid was designated pWPI-Luc2. All constructs were verified by DNA sequencing.

### Cell maintenance, transfection, virion production assays, and protein analyses

293T and HeLa cells were maintained under standard conditions. 293T cells (2.5 x 10^5^) were cotransfected with either 1 μg of either pNL-Luc2-E^-^R^+^, pNL-Luc2-IN/HiBiT, or pNL-Luc2-E^-^-IN/HiBiT-Fin using FuGENE6 (Promega) according to the manufacturer’s instructions. Sixteen hours later, the cells were washed with phosphate-buffered saline (PBS), and then 1 ml of fresh complete medium was added. After 24 hours, the supernatants were harvested, and subjected to an HIV-1 p24-antigen capture enzyme-linked immunosorbent assay (ELISA; XpressBio) to determine the level of virion production. Cells were separated and lysed in 75 μl of cell culture lysis reagent (Promega) or in 75 μl of RIPA buffer (50 mM Tris-HCl, pH 7.4, 150 mM NaCl, 0.5% sodium deoxycholate, 1% Nonidet P-40, 0.1% SDS, and Complete Protease Inhibitor Mixture (Roche Applied Science)). The former cell lysates were subjected to a luciferase assay to normalize the transfection efficiency using a firefly Luciferase Assay System (Promega) with a Centro LB960 luminometer (Berthold). The latter lysates were subjected to Western blot analysis using an anti-p24 monoclonal antibody (1:1,000; Nu24 (16)) an anti-Vif peptide polyclonal antibody (1:200; MBL), and an anti-β-actin mouse monoclonal antibody (1:5,000; Sigma-Aldrich, A5316). Reacted proteins were visualized by chemiluminescence using an ECL Western blotting detection system (GE Healthcare) and monitored using a LAS-3000 imaging system (FujiFilm).

### Virus preparation and infectivity assays

For NL4-3-based virus production, 293T cells (2.5 x 10^5^) were cotransfected with 1 μg of either pNL-Luc2-E^-^R^+^, pNL-Luc2-IN/HiBiT, or pNL-Luc2-E^-^-IN/HiBiT-Fin, and 20 ng of pC-VSVg, with or without 10 ng of pC-hA3G-HA by using FuGENE6. For lentiviral production, 293T cells were cotransfected with 475 ng of either psPAX2 or psPAX2-IN/HiBiT, 475 ng of pWPI-Luc2, and 50 ng of pC-VSVg by using FuGENE6. Sixteen hours later, the cells were washed with PBS, and then 1 ml of fresh complete medium was added. After 24 hours, the supernatants were harvested and treated with 37.5 U/ml DNase I (Roche) at 37°C for 30 minutes. The viral supernatants were subjected to HIV-1 p24 ELISA to measure the p24 antigen. To determine infectivity, HeLa cells (1 x 10^4^) were incubated with the p24 antigen (1 ng) of each viral supernatant. After 48 hours, cells were lysed in 100 μl of lysis buffer, and firefly luciferase activities were determined as described above.

### Primary cell culture

Experiments using human samples were approved by the Medical Research Ethics Committee of the National Institute of Infectious Diseases, Japan. Peripheral blood mononuclear cells (PBMCs) were obtained from healthy volunteer donors who signed an informed consent form. Briefly, PBMCs were isolated by Ficoll-Hypaque gradient centrifugation. The CD4-positive T lymphocytes were purified using a Dynabeads CD4-positive isolation kit (Invitrogen) to obtain CD4-positive T cells, and purified cells were cultured in the presence of 3 μg/ml PHA (Sigma-Aldrich) and 10 U/ml IL-2 (Peprotech) for 72 hours.

### Virus replication assays

CD4-positive T cells (2.5 x 10^5^) were infected for 3 hours with either NL4-3 or NL-IN/HiBiT-Fin virus (25 ng of p24 antigen), washed extensively with serum-free medium, and then cultured in fresh complete medium. Supernatants were sampled every 3 days, and p24 antigen production was quantified by ELISA.

### HiBiT luciferase assays

A high-titer virus stock as a standard sample was prepared by propagating IN/HiBiT-Fin virus in infected M8166 cells and measuring p24 by ELISA. After determining the levels of p24 antigen, the viral supernatant was aliquoted and stored for future use in different HiBiT-based assays, to avoid repeated freeze-thaw cycles. To create a standard curve, the viral stock with known levels of p24 antigen was serially diluted. Either the standards or the samples of interest (25 μl) and LgBiT Protein (1:100)/HiBiT Lytic Substrate (1:50) in Nano-Glo HiBiT Lytic Buffer (25 μl) (Nano-Glo HiBiT Lytic Detection System; Promega) were mixed and incubated for ten minutes at room temperature according to the modified manufacturer’s instructions. Luciferase activity was determined with a Centro LB960 luminometer.

### Transmission electron microscopy

Cells infected with either the WT NL4-3 or NL-IN/HiBiT-Fin virus were harvested after 24 hours with a cell scraper and washed twice with ice-cold PBS. Cells were then prefixed with 2.5% glutaraldehyde and 2% paraformaldehyde in 0.1 M phosphate buffer, pH 7.4, for 2 hours at room temperature, postfixed in 1% osmium tetroxide, and embedded in Epon 812 (TAAB Laboratories). Ultrathin sections were stained with uranyl acetate and lead citrate and then observed under a transmission electron microscope (HT7700; Hitachi) at 80 kV.

### Statistical analyses

Column graphs that combine bars and individual data points were created with GraphPad Prism version 8.04, and coefficient of determination (R2) and regression equations were also calculated with GraphPad Prism.

## ACKNOWLEDGMENTS

We thank D. Trono (Ecole Polytechnique Fédérale de Lausanne, Switzerland) for providing psPAX2 and pWPI. This work was supported by a grant from the Ministry of Health, Labor and Welfare of Japan. The authors have no conflicting financial interests.

## Notes

### Competing Interest Statement

The authors have declared no competing interest.

## REFERENCES

1. Adachi A, Gendelman HE, Koenig S, Folks T, Willey R, Rabson A, Martin MA. 1986. Production of acquired immunodeficiency syndrome-associated retrovirus in human and nonhuman cells transfected with an infectious molecular clone. J Virol 59:284–91.

2. Barre-Sinoussi F, Chermann J, Rey F, Nugeyre M, Chamaret S, Gruest J, Dauguet C, Axler-Blin C, Vezinet-Brun F, Rouzioux C, Rozenbaum W, Montagnier L. 1983. Isolation of a T-lymphotropic retrovirus from a patient at risk for acquired immune deficiency syndrome (AIDS). Science 220:868–871.

3. Goudsmit J, Lange JM, Paul DA, Dawson GJ. 1987. Antigenemia and antibody titers to core and envelope antigens in AIDS, AIDS-related complex, and subclinical human immunodeficiency virus infection. J Infect Dis 155:558–60.

4. Dixon AS, Schwinn MK, Hall MP, Zimmerman K, Otto P, Lubben TH, Butler BL, Binkowski BF, Machleidt T, Kirkland TA, Wood MG, Eggers CT, Encell LP, Wood KV. 2016. NanoLuc Complementation Reporter Optimized for Accurate Measurement of Protein Interactions in Cells. ACS Chem Biol 11:400–8.

5. Vermeire J, Naessens E, Vanderstraeten H, Landi A, Iannucci V, Van Nuffel A, Taghon T, Pizzato M, Verhasselt B. 2012. Quantification of reverse transcriptase activity by real-time PCR as a fast and accurate method for titration of HIV, lenti- and retroviral vectors. PLoS One 7:e50859.

6. Pizzato M, Erlwein O, Bonsall D, Kaye S, Muir D, McClure MO. 2009. A one-step SYBR Green I-based product-enhanced reverse transcriptase assay for the quantitation of retroviruses in cell culture supernatants. J Virol Methods 156:1–7.

7. Dang Y, Wang X, Zhou T, York IA, Zheng YH. 2009. Identification of a novel WxSLVK motif in the N terminus of human immunodeficiency virus and simian immunodeficiency virus Vif that is critical for APOBEC3G and APOBEC3F neutralization. J Virol 83:8544–52.

8. Yamashita T, Kamada K, Hatcho K, Adachi A, Nomaguchi M. 2008. Identification of amino acid residues in HIV-1 Vif critical for binding and exclusion of APOBEC3G/F. Microbes Infect 10:1142–9.

9. Russell RA, Pathak VK. 2007. Identification of two distinct human immunodeficiency virus type 1 Vif determinants critical for interactions with human APOBEC3G and APOBEC3F. J Virol 81:8201–10.

10. Tian C, Yu X, Zhang W, Wang T, Xu R, Yu XF. 2006. Differential requirement for conserved tryptophans in human immunodeficiency virus type 1 Vif for the selective suppression of APOBEC3G and APOBEC3F. J Virol 80:3112–5.

11. Delenda C, Gaillard C. 2005. Real-time quantitative PCR for the design of lentiviral vector analytical assays. Gene Ther 12 Suppl 1:S36–50.

12. Geraerts M, Willems S, Baekelandt V, Debyser Z, Gijsbers R. 2006. Comparison of lentiviral vector titration methods. BMC Biotechnol 6:34.

13. Tada T, Zhang Y, Koyama T, Tobiume M, Tsunetsugu-Yokota Y, Yamaoka S, Fujita H, Tokunaga K. 2015. MARCH8 inhibits HIV-1 infection by reducing virion incorporation of envelope glycoproteins. Nat Med 21:1502–7.

14. Kinomoto M, Kanno T, Shimura M, Ishizaka Y, Kojima A, Kurata T, Sata T, Tokunaga K. 2007 All APOBEC3 family proteins differentially inhibit LINE-1 retrotransposition. Nucleic Acids Res 35:2955–64.

15. Wiznerowicz M, Trono D. 2003. Conditional suppression of cellular genes: lentivirus vector-mediated drug-inducible RNA interference. J Virol 77:8957–61.

16. Tsunetsugu-Yokota Y, Akagawa K, Kimoto H, Suzuki K, Iwasaki M, Yasuda S, Hausser G, Hultgren C, Meyerhans A, Takemori T. 1995. Monocyte-derived cultured dendritic cells are susceptible to human immunodeficiency virus infection and transmit virus to resting T cells in the process of nominal antigen presentation. J Virol 69:4544–7.

